# Comparison of cerebrospinal fluid biomarkers relevant to neurodegenerative diseases in healthy cynomolgus and rhesus macaque monkeys

**DOI:** 10.1101/2021.03.01.433384

**Authors:** Emma L. Robertson, Susan E. Boehnke, Natalia M. Lyra e Silva, Brittney Armitage-Brown, Andrew Winterborn, D.J. Cook, Fernanda G. De Felice, Douglas P. Munoz

## Abstract

**INTRODUCTION:** Non-human primates are important translational models of neurodegenerative disease. We characterized how species, sex, age, and site of sampling affected concentrations of key biomarkers of neurodegeneration.

**METHODS:** Amyloid-beta (Aβ40, Aβ42), tau (tTau, pTau), and neurofilament light (NFL) in CSF were measured in 82 laboratory-housed naïve cynomolgus and rhesus macaques of both sexes.

**RESULTS:** Aβ40, Aβ42, and NFL were significantly higher in rhesus compared with cynomolgus macaques. tTau and NFL were higher in males. pTau was not affected by species or sex. Site of acquisition only affected NFL, with NFL being higher in CSF acquired from lumbar compared with cisterna magna puncture.

**DISCUSSION:** Normative values for key neurodegeneration biomarkers were established for laboratory housed cynomolgus and rhesus macaque monkeys. Differences were observed as a function of species, sex and site of CSF acquisition that should be considered when employing primate models.

**Research In Context:**

1. Systematic review: We reviewed reports characterizing CSF biomarkers of neurodegenerative diseases in non-human primates – an increasingly important model of disease - revealing that studies with laboratory housed macaque monkeys were of small sample size, with a paucity of data about how biomarkers varied as a function of species, sex, age, and site of acquisition.
2. Interpretation: To address this gap, we collected CSF from 82 naïve laboratory housed male and female macaques of two species and measured Aβ40, Aβ42, tTau, pTau, and NFL. In addition to providing normative statistics for concentrations of these biomarkers, we revealed various species and sex differences.
3. Future directions: Establishing normative values of biomarkers is an important step to the efficient development of cynomolgus and rhesus macaques as models of neurodegenerative disorders such as Alzheimer’s disease. Reference values reduce the need for large control groups by which to compare with disease model animals.

## Introduction

Cerebrospinal fluid (CSF) biomarkers are increasingly used to diagnose and track progression and evaluate treatment of various human neurological disorders. For example, core CSF biomarkers involved with Alzheimer’s disease (AD) include amyloid-beta 1-40 (Aβ40), amyloid-beta 1-42 (Aβ42), total tau (tTau), and phosphorylated tau (pTau)^1,2^. Additionally, neurofilament light (NFL), a protein providing structural support to the neural cytoskeleton, has been identified as a marker of general axonal degradation and is also being investigated in a variety of neurodegenerative disorders such as AD^3–7^, Parkinson’s disease^5,8,9^, multiple sclerosis^10–13^, and Huntington’s disease^3,14^.

Old-world monkeys, such as cynomolgus (*Macaca fascicularis*) and rhesus macaques (*Macaca mulatta*), are important models for human neurological disorders due to their similarities in brain architecture, anatomy, physiology, and behaviour^15,16^. As many neurodegenerative disorders involve impairment of higher cognitive functions, monkeys have the potential to advance our knowledge of these disorders and validate treatments. To fully validate monkey models of neurodegenerative disorders, it is important to better understand the factors that affect these CSF biomarkers. There are several non-human primate (NHP) models of AD and aging being developed^17–25^, of which only a few characterize the changes and progression of CSF biomarkers^17–20,22,24,25^. Natural aging models in NHPs have shown development of amyloid plaque deposition and phosphorylated tau accumulation in the brain parenchyma^22,26–28^, with some showing age-associated changes in CSF biomarkers^22,26^.

Lumbar punctures (LPs) can be performed at the intervertebral spaces in NHPs to collect CSF samples, aiding in the translational aspect to human clinical work, which typically employs LPs. However, it is also common for CSF to be obtained from cisterna magna puncture in NHPs. Therefore, it is important to contrast and assess differences in CSF samples acquired from these two sites. For example, we have previously demonstrated that LPs, but not cisterna manga punctures, significantly elevate NFL for several weeks^29^.

Despite significant progress in development of NHP models of disease, little is known about the range of normative CSF values of Aβ, tau, and NFL in healthy NHPs in laboratory housing conditions. Given the limited samples sizes of many primate studies, it is difficult to interpret pathological CSF changes in these models without having references ranges with which to compare. It is also unknown how factors such as species, sex, age, and location of the sample taken affect biomarker levels in NHPs, independent of any neurological disease. To address these gaps in knowledge, we characterized normative values for Aβ40, Aβ42, tTau, pTau, and NFL in a colony of naïve cynomolgus and rhesus macaques of both sexes, with CSF acquired via LP or cisterna magna puncture.

## Methods

### Subjects

All NHPs were housed at the Centre for Neuroscience Studies at Queen’s University (Kingston, Ontario, Canada) under the care of a lab animal technician and the Institute Veterinarian. All procedures were approved by the Queen’s University Animal Care Committee and were in full compliance with the Canadian Council on Animal Care (Animal Care Protocol Munoz, 2011-039-Or).

LPs were performed on a total of 82 animals (Fig. 1): 19 male and 30 female rhesus macaques (*Macaca mulatta*, ages: 3-12 years, body weight: 5.4-18 kg) and 28 male and 5 female cynomolgus macaques (*Macaca fascicularis*, ages: 2-9 years, body weight: 3.4-10.7 kg). Animals were housed indoors either in small groups (n=49) or individually (n=33), and kept on a 12:12-hr light:dark cycle starting at 7am. They were fed a standard diet of high-protein or high-fiber monkey chow and supplemented with fresh fruit and vegetables. On CSF collection day, animals were fasted but had access to water ad libitum. Daily enrichment was provided through foraging, puzzle toys, swings, ropes, perches, mirrors, etc. All animals were experimentally naïve when the CSF samples were obtained except for 3 animals for whom we were unable to acquire CSF on the first attempt. On a separate day, a second attempt was required to obtain a sample.

**Figure 1.**
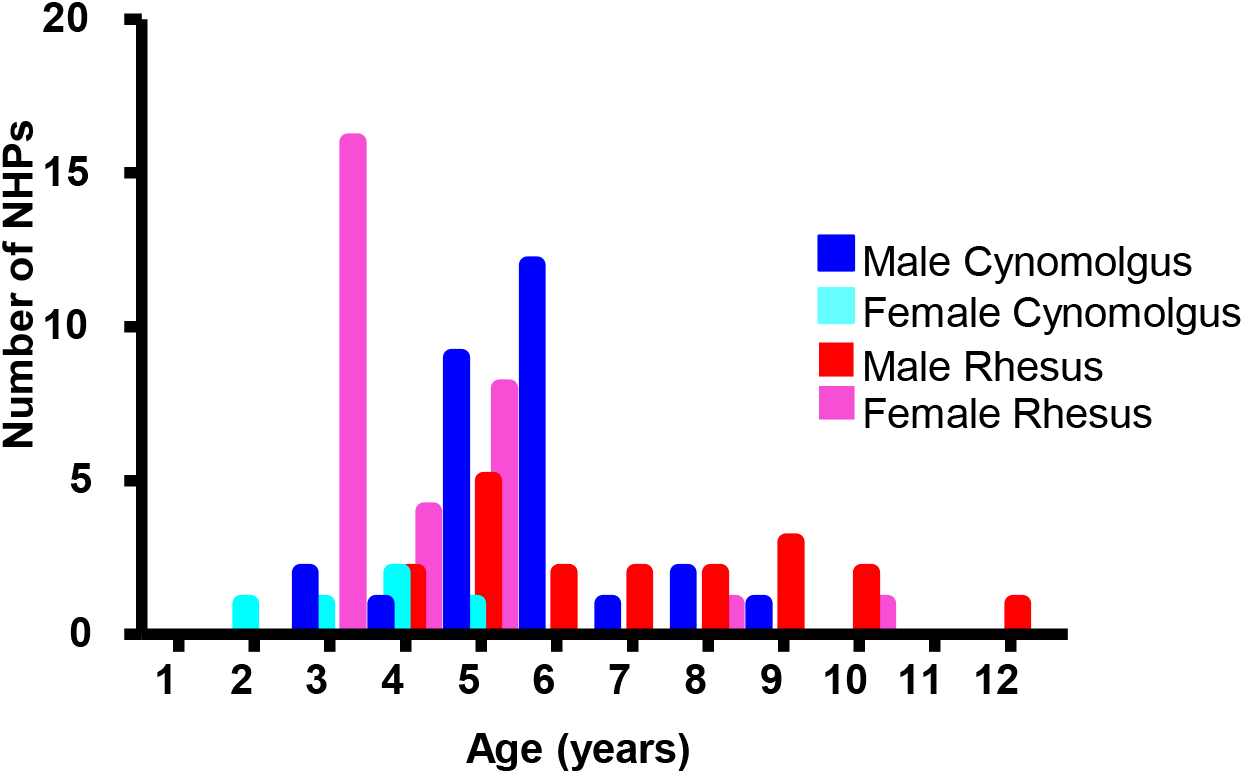
Demographics of non-human primate colony. CSF from 82 animals was analyzed: 28 male and 5 female cynomolgus macaques (*Macaca fascicularis*, ages: 2-9 years, body weight: 3.4-10.7 kg) and 19 male and 30 female rhesus macaques (*Macaca mulatta*, ages: 3-12 years, body weight: 5.4-18 kg) (Fig. 2.1). All animals were experimentally naïve.

Cisterna magna punctures were performed in a subset of 16 of the 82 for which CSF was acquired by LP (7 male and 3 female cynomolgus macaques (ages: 2-6 years, body weight: 3.4-10.7 kg) and 6 male rhesus macaques (ages: 4-9 years, body weight: 5.4-18 kg). All 16 samples were analyzed for NFL and 11 samples were analyzed for Aβ40, Aβ42, tTau, and pTau. The cisterna puncture was performed on a separate occasion at least 5 months after the LP was performed.

### CSF Collection and Storage

CSF was collected as previously described^29^. A trained veterinarian or veterinary technician performed LPs and cisterna magna punctures between 9am-1pm. Animals were sedated with ketamine (5-15 mg/kg, intramuscular) and masked briefly, only when required, with isoflurane (1-3%) and oxygen (2%) to minimize movement. If additional procedures were planned (e.g. MRI, surgery), anaesthesia was induced with ketamine (10 mg/kg, intramuscular) and diazepam (5 mg/kg, intramuscular). Glycopyrrolate (0.013 mg/kg, intramuscular) was given and the animal was intubated. Anaesthesia was maintained using isoflurane (1-3%) and oxygen (2%). In these relatively rare cases (n=3), CSF sampling was the first procedure conducted to minimize any anesthetic effects.

For LP sampling, animals were placed in lateral recumbency and the superior iliac crest was palpated. The lumbar area was shaved and cleaned using chlorhexidine, alcohol, and betadine. Depending on animal size, a 20g or 22g Quincke spinal needle (BD™) was inserted into the intrathecal space between L4/5, or in some cases, L3/4 or L5/6. CSF was allowed to drip by gravity into a sterile 1.5ml polypropylene Eppendorf tube (Axygen) and then immediately placed on ice.

For cisterna magna sampling, animals were placed in lateral or sternal recumbency and the area between the occipital protuberance to the third cervical vertebrae was shaved and cleaned. A 23g needle connected to a 1ml (BD Luer-Lok) polypropylene syringe was inserted into the midline of the neck, with the tip of the needle pointing towards the nose^30^. No adverse neurological effects were observed following LP or cisterna magna punctures.

If the CSF sample was visibly contaminated with blood, it was centrifuged at 1800g for 10 minutes at 4°C to remove blood cells. Samples were stored as 120ul aliquots within 30 minutes of collection in 0.6 ml sterile polypropylene tubes (Axygen) at -80°C. Prior to biochemical analysis, samples did not undergo any freeze-thaw cycles.

## CSF Biomarker analysis

CSF samples were thawed in a biological safety cabinet just before analysis. To measure Aβ40, Aβ42, pTau (pThr181) and tTau, a MILLIPLEX Human Amyloid Beta Tau Magnetic Bead Panel (HNABTMAG-68K, EMD Millipore, Billerica, MA, USA) was completed on a Biorad Luminex platform according to the manufacturer’s instructions. NFL was measured using a commercial sandwich ELISA (NF-light ELISA kit, UmanDiagnostics, Umeå, Sweden) performed according to the manufacturer’s instructions. Within plate and interplate coefficients of variation were <15% for the multiplex; and <10% for NFL respectively.

### Exclusion Criteria

Exclusion criteria included: 1. Any values below the detection limits; 2. For NFL analysis, animals that received a recent LP, as this procedure was shown to elevate NFL (but not other Aβ/tau biomarkers) for several weeks^29^; 3. Any values that were more than four standard deviations above the mean and flagged by Grubb’s test as outliers (*p* <.01). For CSF collected via LP, 1 female rhesus was removed from Aβ40/Aβ42. 1 male cynomolgus, 1 male rhesus, and 5 female rhesus macaques were removed from tTau. One male cynomolgus and 3 male rhesus macaques were removed from NFL. For CSF collected via cisterna magna, 1 male and female cynomolgus macaque was removed from tTau, 1 male cynomolgus macaque was removed from pTau and another was removed from NFL.

### Data/Statistical Analysis

Statistical analyses were performed with SPSS version 26. Pearson correlations were used to test for relationships between age and biomarker levels. To examine the effects of species and sex on Aβ40,

Aβ42, tTau, pTau, and NFL while controlling for age, a two-way ANCOVA was conducted. To compare location differences of LP vs cisterna magna puncture, the Mann-Whitney U test for independent samples was used because the data was not normally distributed. A Pearson correlation was used to assess the association between each biomarker. *P* values <0.05 were considered significant.

## Results

A two-way ANCOVA was conducted on each of the biomarkers to assess the role of species and sex. The age range was small, and significant main effects of age were only observed for tTau, thus we removed age effects by adding age as a covariate in the analysis. The means, age-adjusted means, standard deviations, and standard errors of each biomarker are presented in Table 1. The raw values for each animal are plotted separated by species and sex for each biomarker in Figure 2A-E.

**Table 1.** **Means, Age-Adjusted Means, Standard Deviations, and Standard Errors for Aβ40, Aβ42, tTau, pTau, and NFL concentration in cynomolgus and rhesus macaques**

|  |  | Cynomolgus |  | Rhesus |  |
| --- | --- | --- | --- | --- | --- |
|  |  | Male | Female | Male | Female |
| Aβ40 pg/ml | *N* | 28 | 5 | 19 | 29 |
|  | *M* | 1779.27 | 1539.33 | 2271.48 | 3083.87 |
|  | (*SD*) | (721.64) | (585.17) | (774.97) | (1006.50) |
|  | *M*<sub>adj</sub> | 1813.91 | 1391.52 | 2426.73 | 2974.20 |
|  | (*SE*) | (159.18) | (384.73) | (215.35) | (169.80) |
| Aβ42 pg/ml | *N* | 28 | 5 | 19 | 29 |
|  | *M* | 317.29 | 266.68 | 397.60 | 608.89 |
|  | (*SD*) | (125.17) | (141.25) | (143.89) | (274.60) |
|  | *M*<sub>adj</sub> | 318.27 | 262.50 | 401.99 | 605.78 |
|  | (*SE*) | (37.82) | (91.41) | (51.17) | (40.37) |
| tTau pg/ml | *N* | 27 | 5 | 18 | 25 |
|  | *M* | 286.71 | 199.37 | 310.71 | 237.03 |
|  | (*SD*) | (114.28) | (70.09) | (162.23) | (81.99) |
|  | *M*<sub>adj</sub> | 293.85 | 156.23 | 355.69 | 205.56 |
|  | (*SE*) | (21.24) | (50.90) | (29.42) | (24.04) |
| pTau pg/ml | *N* | 28 | 5 | 19 | 30 |
|  | *M* | 40.93 | 33.43 | 39.87 | 33.67 |
|  | (*SD*) | (17.36) | (14.21) | (17.82) | (9.47) |
|  | *M*<sub>adj</sub> | 41.29 | 31.94 | 41.439 | 32.60 |
|  | (*SE*) | (2.84) | (6.87) | (3.85) | (2.98) |
| NFL pg/ml | *N* | 27 | 5 | 16 | 30 |
|  | *M* | 372.22 | 184.67 | 505.42 | 443.19 |
|  | (*SD*) | (142.15) | (58.39) | (231.01) | (251.74) |
|  | *M*<sub>adj</sub> | 375.17 | 175.87 | 515.08 | 436.96 |
|  | (*SE*) | (40.83) | (96.21) | (58.30) | (41.62) |

Abbreviations: Aβ , amyloid beta; CSF, cerebrospinal fluid; NFL, neurofilament light; pTau, phosphorylated tau; SD, standard deviation; SE, standard error; tTau, total tau.

**Figure 2.**
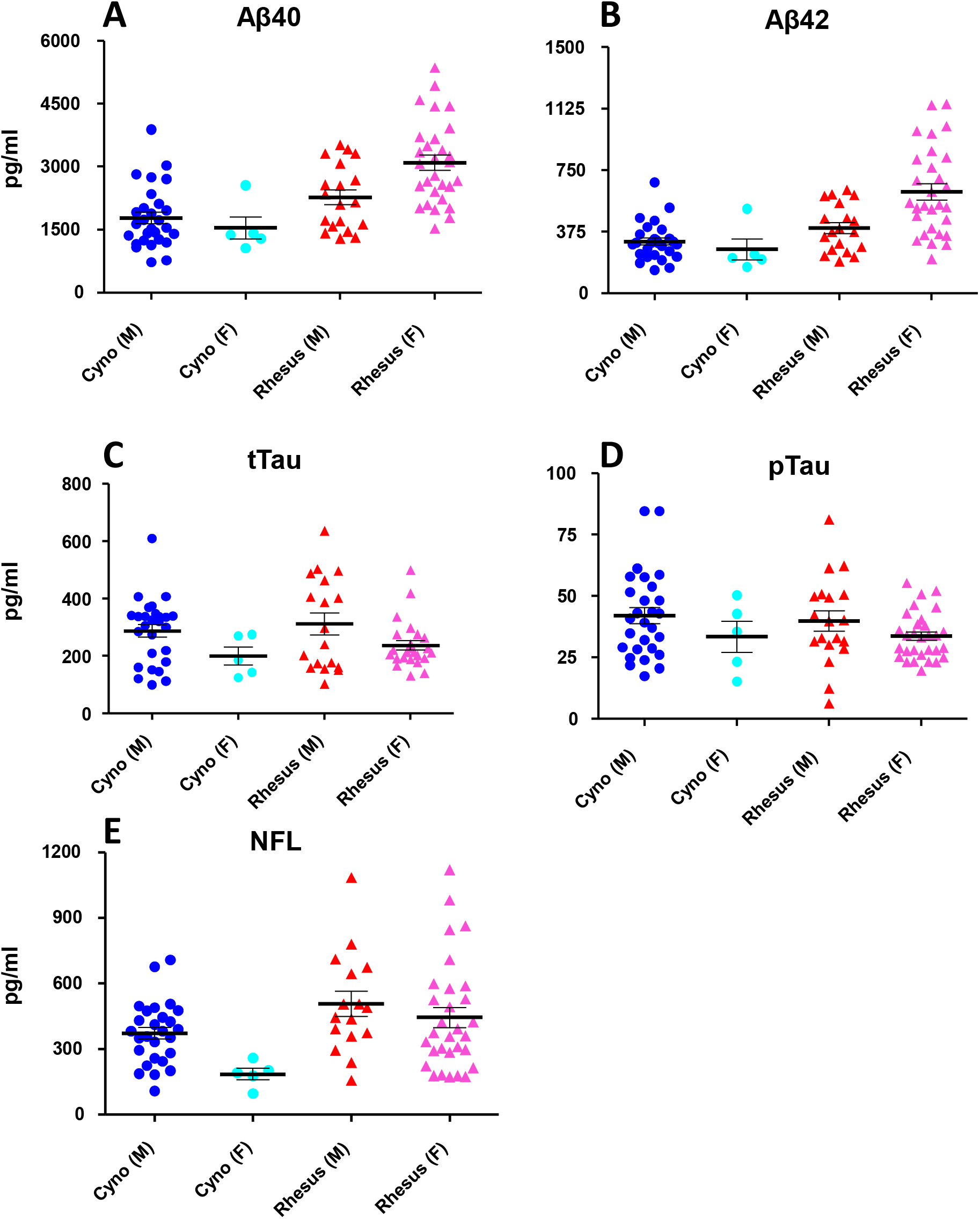
Species and sex comparison for CSF Aβ, tau, and NFL biomarkers. A) Aβ40 concentrations were higher in Rhesus macaques, and the species difference was larger for females. B) Aβ42 concentrations were higher in Rhesus macaques, and the species difference was larger for females. C) tTau concentrations were higher for males. D) pTau concentrations did not differ between groups (*p*’s >0.05) E) NFL concentrations were higher in Rhesus macaques, and males were higher than females.

### Aβ40 and AB42

For both Aβ40 and AB42, there was a main effect of species [Aβ40: F(1,76)=20.53, p<0.001, η2 = 0.21; Aβ42: F(1,76)=13.748, p<0.001, η2=0.15] with rhesus macaques [Aβ40: 2700.46 pg/ml; Aβ42: 503.89 pg/m] having higher concentrations than cynomolgus macaques [Aβ40: 1602.72 pg/ml; Aβ42: 290.43 pg/ml]. There was no main effect of sex in either biomarker (p=0.82; p=0.26) but there was a statistically significant interaction between species and sex [Aβ40: F(1,76) = 4.14, p =0.045, partial η2 = 0.052; Aβ42: F(1,76)=5.252, p=0.025, η2=0.065]. Analysis of simple effects indicated that the effect of species was greater for females [Aβ40: rhesus 2974.2 pg/ml vs cynomolgus 1391.52 pg/ml; p<0.001, Aβ42: rhesus 605.78 pg/ml vs cynomolgus 262.50 pg/ml; p<0.001] than males (Aβ40 rhesus 2426.73 pg/ml vs cynomolgus 1813.91 pg/ml; p=0.02, Aβ42: rhesus 401.99 pg/ml vs cynomolgus 318.37 pg/ml; p=0.179).

### tTau and pTau

For tTau, there was a main effect of sex [F(1,70)=14.88, p<0.001, η2 = 0.175], with males (324.77 pg/ml) having higher concentrations than females (180.89 pg/ml). There was not a significant main effect of species [F(1,70)=2.911, p=0.092, η2 = 0.040] or an interaction [F(1,70)=0.038, p=0.845, η2=0.001]. For pTau there was no main effect of species or sex and there was no interaction (all p’s >0.05).

### NFL

For NFL, there was a main effect of species [F(1,73)=10.64, p=0.002, η2=0.127] with rhesus (476.02 pg/ml) having higher concentrations than cynomolgus (275.52 pg/ml). There was also a main effect of sex [F(1,73)=3.85, p=0.05, η2=0.050] with males(445.12 pg/ml) having higher values than females (306.42 pg/ml). There was not an interaction [F(1,73)=1.006, p=0.319, η2 = 0.014].

### Effects of age

We wanted to determine if any of the biomarkers changed across the relatively narrow range of ages (2-12 yrs) represented within the colony (Fig.1). Collapsed across species and sex, there was a negative correlation in Aβ40 and Aβ42 with age (*R*^2^ = 0.065, *p* = .021; *R*^2^ = 0.092, *p* =0.006), but no significant correlation of tTau, pTau or NFL with age. When separated by species and sex, Aβ40 and Aβ42 in female rhesus macaques decreased significantly with age (*R*^2^ = 0.173, *p* = 0.025; *R*^2^ = 0.151, *p* =0.037), and tTau and pTau concentration decreased significantly with age in male rhesus macaques (*R*^2^= 0.427, *p* =0.003; *R*^2^ = 0.208, *p* = 0.05). No other biomarkers were correlated with age (*p*’s >0.05).

### Effects of CSF location

In a subset of the NHPs (n=16) for whom CSF had been obtained by LP, CSF was collected through the cisterna magna on a separate occasion (see Methods). Only NFL differed between sample sites - it was significantly lower in cisterna samples (Mann Whitney U = 45.5, z = -2.779, *p* =0.004) (Fig. 3). The median percent change ((lumbar puncture – cisterna puncture)/cisterna puncture*100) in NFL was -42% (IQR=51%) (Fig. 3F). That NFL is significantly lower in cisterna samples was previously reported

**Figure 3.**
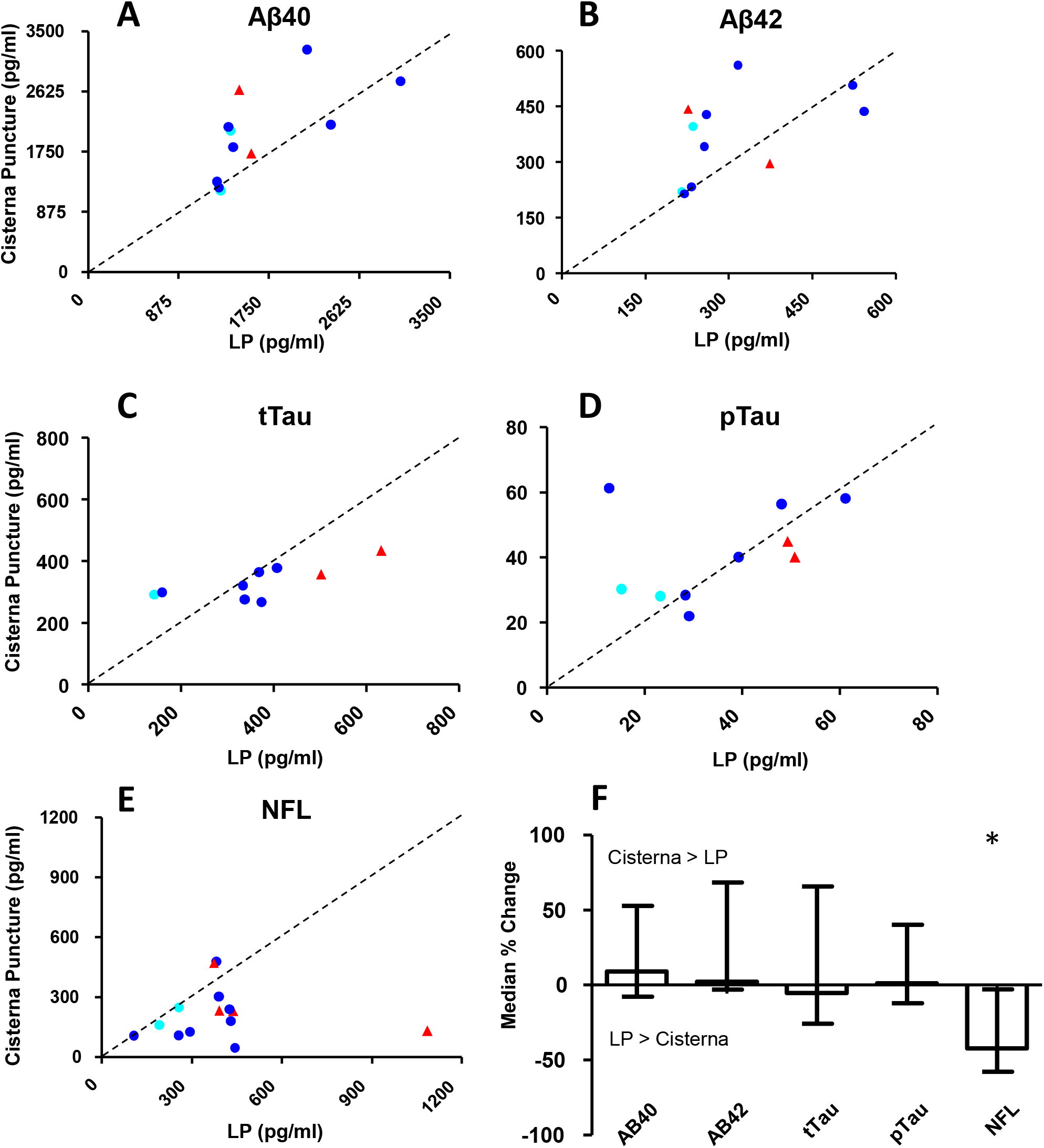
Comparison of biomarkers taken from NHPs that had both lumbar punctures and cisterna magna punctures. A subset of NHPs from the colony had both lumbar and cisterna punctures are values from each animal are plotted for each biomarker: A) Aβ40 B) Aβ42 C) tTau D) pTau E) NFL. F) The median percent change ((lumbar puncture – cisterna puncture)/cisterna puncture*100) is plotted for each biomarker along with the interquartile range. NFL in CSF taken from the cisterna magna was significantly lower than from CSF taken from the lumbar area (Mann Whitney U = 45.5, z = -2.779, p = .004). All other p’s >.05

### Amyloid-beta and tau biomarkers are positively correlated with each other

Aβ40 and Aβ42 were positively correlated with each other [F(1,79) = 752.559, *p* <0.001, r(79) = 0.951] (Fig. 2.5). tTau and pTau were also positively correlated [F(1,73) = 20.622, *p* <0.001, r(73) = 0.469] (Fig. 4). No other combination of biomarker correlations reached significance (Data not shown, all *p*’s >0.09).

**Figure 4.**
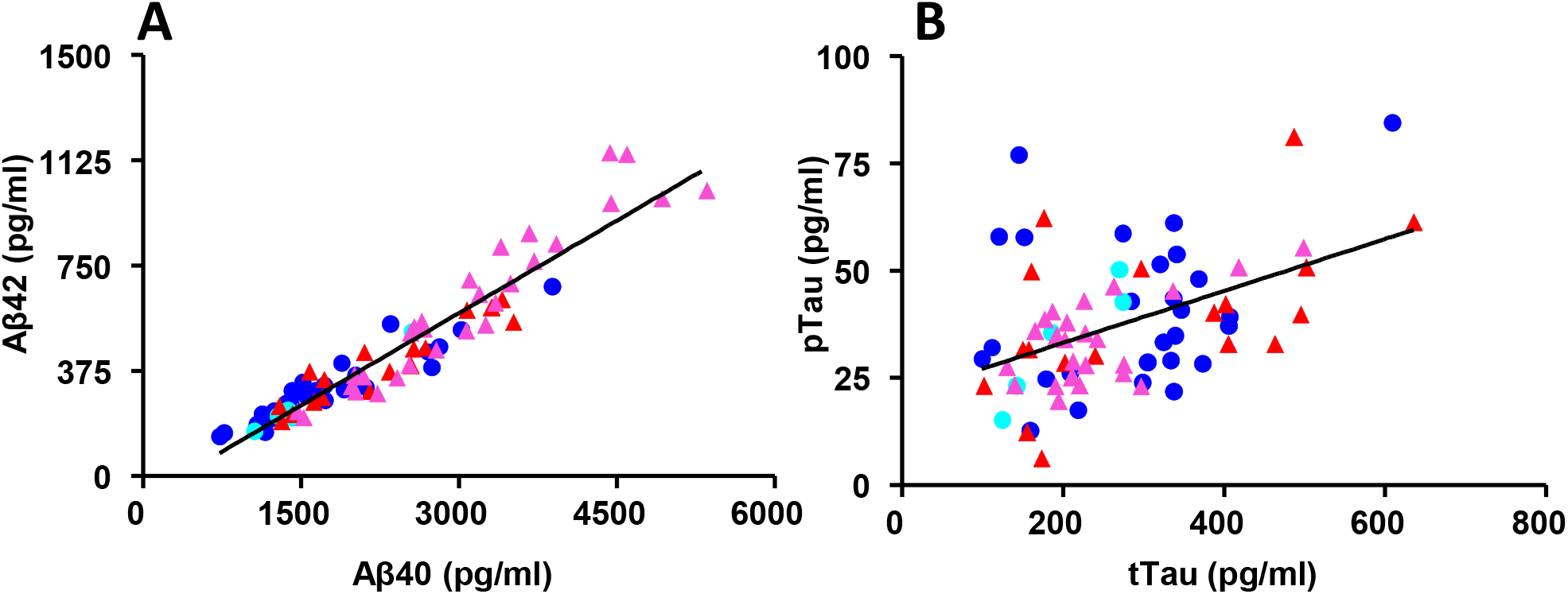
Amyloid-beta and tau biomarkers are correlated with each other. Each point represents one animal. Aβ40 and Aβ42 show a significant positive correlation. Linear regression: F(1,79) = 752.56, p <0.001, r(79) = 0.95. tTau and pTau also show a positive correlation. Linear regression: F(1,73) = 20.62, p < 0.001, r(73) = 0.47. No other biomarker combinations were correlated(all p’s >0.05.).

## Discussion

Using a laboratory housed colony of cynomolgus and rhesus macaques, we investigated the effects of species, sex, age, and location of sample acquisition on common CSF biomarkers used to characterize neurodegenerative diseases such as AD. These data help establish normative values for these biomarkers in two commonly used species of macaque and will be important reference points for efficient use and development of primate models of neurodegenerative disease.

In summary, concentrations of Aβ40, Aβ42, and NFL were all significantly higher in rhesus macaques compared to cynomolgus macaques. tTau and NFL concentrations were significantly higher in males compared to females. There were interactions between species and sex for both Aβ40 and Aβ42, with the species difference being greater for females than for males. For pTau there was no effect of species or sex. As would be expected, strong correlations were observed between Aβ40 and Aβ42 and between tTau and pTau, but we observed no other correlations between biomarkers. As age increased, Aβ biomarkers decreased in female rhesus; and tau biomarkers decreased in male rhesus. Finally, CSF samples from the cisterna magna puncture were lower in NFL when compared to CSF obtained from LP. No other biomarker was affected by sampling location.

## Relation to previous NHP studies

### Rhesus Macaques

Zhao et al. collected CSF via LP in young (n=5), mature (n=3) and old (n=4) rhesus macaques and observed Aβ40 values ranging from about 1700-2300 pg/ml and Aβ42 values ranging from about 75-125 pg/ml ^25^. Our observed values were generally higher for both biomarkers, especially Aβ42. This could be due to differences in assays, as we used a Luminex multiplex (Millipore, USA) while they used individual ELISA kits for each biomarker (ImmunoBiological Laboratories, Japan). In another study done by Li et al., CSF was obtained by LP and Aβ40 and Aβ42 were measured in young (n = 15), middle-aged (n = 22), and aged (n = 19) groups^31^. Their reported values were quite similar to ours, and Aβ40 values ranged from approximately 1000-3300 pg/ml and Aβ42 ranged from 100-800 pg/ml^31^.

Beckman et al. looked at several of the same biomarkers we report here in a group of female rhesus (n=6) that were undergoing intracerebroventricular injections of Aβ-Oligomers^18^ to induce a monkey model of AD^21^. They obtained CSF via cisterna puncture, and found baseline values for Aβ40 that were slightly higher than ours (∼3500-4200 pg/ml), and for Aβ42 they were somewhat lower than ours (∼100-125 pg/ml). Their values for tTau (200-225pg/ml) were similar to ours. For pTau, they examined different phosphorylation sites (pS199, pS231, pS396) than we did (pThr181), making comparison difficult. Their values for NFL were an order of magnitude higher (3000-4000pg/ml) than we observed suggestive of axonal damage in their animals, probably due to the implant surgery those animals had undergone to allow for the intracerebroventricular injections. We could find no other reference values in the literature for tTau, pTau, or NFL in CSF of naïve rhesus macaques.

### Cynomolgus Macaques

In a study done in male and female cynomolgus macaques, young animals (n = 12) and aged animals (n = 20) had tTau values averaging 639 pg/ml and 544 pg/ml, respectively^20^. These values were on the high end of our reported range. Additionally, pTau (pS396) was reported to average 50.4 pg/ml and 43.7 pg/ml, respectively, which was very similar to our data, despite our analysis of pTau (threonine 181 instead of serine 396). In another study by Darusman et al.^19^, 12 aged cynomolgus macaques (male and female) were grouped into low-performers or high-performers in a behavioural task. They found Aβ42 values averaging 258.8 pg/ml and 454.7 pg/ml, in line with our values. Additionally, tTau was found to be 496.4 pg/ml and 227.8 pg/ml, values that roughly corresponded with ours. Again, they characterized pTau serine 396 and found it to be 44.1 pg/ml and 29.2 pg/ml.

Finally, in another study involving cynomolgus macaques, Yue et al.^24^ also characterized Aβ40 and Aβ42 in males and females with CSF obtained by LP. Values of Aβ40 ranged from approximately 100 ng/ml – 300 ng/ml and Aβ42 ranged from 2-22 ng/ml. Converted to pg/ml, these values are orders of magnitude higher than our values (100,000 – 300,000 and 2,000 to 22,000 pg/ml), suggestive of a conversion error in their analysis. Assuming their true values were 1000-3000 and 20-220 pg/ml, the values in the same range as ours. pTau values in this cohort were found to be 10-90 pg/ml, which was also similar to the range of values observed in our data. While Yue et al, did not specify the epitope at which Tau was phosphorylated for their test, the INNOTEST ELISA they used tests for phosphorylation of tau at 181, as was done in our study.

### African Green Monkeys

Biomarkers of neurodegeneration have also been tracked in African green monkeys (AGM, *Chlorocebus aethiops sabaeus*), another species of old-world monkey. While not a macaque, this species is increasingly used as a model of aging and neurological disease. Research animals of this species are predominately sourced from an inbred strain that emerged from founder animals imported from Africa to St. Kitts island in the Caribbean in the 17^th^ Century. This makes them desirable because they have less genetic diversity. In four studies, cisterna samples were collected in male and female AGMs^22,26,32,33^. Lemere et al. reported a baseline mean Aβ40 concentration of ∼10,500 pg/ml (n=10) and a mean Aβ42 concentration of ∼1350 pg/ml (n=10)^33^. Cramer et al. analyzed CSF (n=11) and reported Aβ40 values ranging from 4000-9000 pg/ml, Aβ42 ranging 100-300 pg/ml, tTau ranging 500-5000 pg/ml, and pTau ranging 25-120 pg/ml ^26^. Consistently, Latimer et al. analyzed CSF and found Aβ42 ranging from approximately 120-600 pg/ml (n=9) and pTau(serine 181) ranging from 10-40 pg/ml (n=13)^22^. Finally, Chen et al. collected CSF from 329 AGM (0-20+ years old) and characterized Aβ40, Aβ42, Tau, and pTau ^32^. On average, Aβ40 was 10267 pg/ml, Aβ42 was 506 pg/ml, Tau was 23 pg/ml, and pTau was 30 pg/ml. Given their value for Tau was lower than that for pTau, Chen et al. discounted their tTau measurement. Other than for measurement of tTau, all 4 of these papers had relatively similar biomarker values.

In general, AGM monkeys appear to have much higher values of Aβ40, Aβ42 and possibly Tau compared to cynomolgus and rhesus macaques. This comparison suggests that a species difference may exist, in particular for Aβ, in which AGM > Rhesus Macaques > Cynomolgus Macaques. This may reflect differences in assay binding between the species, or true differences in amyloid load. Further research should clarify this.

### Comparison of age and sex effects in the literature

Due to the current colony demographics available to us, CSF samples were collected in NHPs younger than 12 years (Fig 1). While we saw some effect of age in Aβ40 and Aβ42 in rhesus macaques, we had limited animals of older age, and the ranges of ages between male and female cynomolgus and rhesus macaques differed. Therefore, our results should be interpreted with caution. With this caveat, our results indicating that Aβ biomarkers decrease with age in female rhesus is consistent with other published data. A few studies with limited sample sizes have revealed a significant decrease with age in CSF Aβ40 levels in both male and female aged cynomolgus and rhesus macaques^24,31^ and also Aβ42^31^. On the other hand, Zhao et al report no change in Aβ40, but an increase in Aβ42 with aged rhesus macaques^25^. In other studies done in AGMs, CSF Aβ40 was also observed to decrease with age^22,32^. Latimer et al. also reported a significant decrease in Aβ42^22^, while another study showed it trending down^26^.

Furthermore, we also found that both tTau and pTau decrease with age in male rhesus macaques. While we found no correlation with age and tau biomarkers in cynomolgus macaques, this was consistent with other previous reports^20,24^ and also similar to that observed in AGMs^22,32^. However, Cramer et al. reported a 2-fold increase in both tTau and pTau biomarkers with age ^26^. This may indicate a species difference between cynomolgus macaques and AGMs. To better compare our age correlations with other published data, further CSF samples would need to be acquired from macaques older than 12 years.

In conclusion, we characterized Aβ40, Aβ42, tTau, pTau, and NFL biomarkers in CSF of cynomolgus and rhesus macaques as a function of sex and age, providing the largest reference values for laboratory housed animals of these species to date. These reference values will be useful benchmarks by which to compare CSF from primate models of neurological disorders generated using these species.

